# 40 Hz acoustic stimulation decreases amyloid beta and modulates brain rhythms in a mouse model of Alzheimer’s disease

**DOI:** 10.1101/390302

**Authors:** Juho Lee, Seungjun Ryu, Hyun-Ju Kim, Jieun Jung, Boreom Lee, Tae Kim

**Affiliations:** Department of Biomedical Science and Engineering, Gwangju Institute of Science and Technology (GIST), Gwangju, South Korea; Department of Brain Science, University of Ulsan College of Medicine, Seoul, South Korea; School of Life Sciences, GIST, South Korea

**Keywords:** Gamma band oscillations, acoustic stimulation, amyloid beta, microglia, brain connectivity, 5XFAD

## Abstract

**Introduction:** The accumulation of amyloid-beta (Aβ) is one of the neuropathologic hallmarks of Alzheimer’s disease (AD) and abnormal gamma band oscillations and brain connectivity have been observed. Recently, a therapeutic potential of gamma entrainment of the brain was reported by Iaccarino *et al*. However, the affected areas were limited to hippocampus and visual cortex. Therefore, we sought to test the effects of acoustic stimulation in a mouse model of AD.

**Methods:** Freely moving 6-month-old 5XFAD mice with electroencephalogram (EEG) electrodes were treated with daily two-hour acoustic stimulation at 40Hz for 2 weeks. Aβ and microglia were evaluated by immunohistochemistry and ELISA. Evoked and spontaneous gamma power were analyzed by wavelet analysis. Coherence, phase locking value (PLV), and cross-frequency coupling were analyzed.

**Results:** The number of Aβ plaques decreased in the pre-and infralimbic (PIL) and hippocampus regions and soluble Aβ-40 and Aβ-42 peptides in PIL in the acoustic stimulation group. We also found that the number of microglia increased in PIL and hippocampus. In EEG analysis, evoked gamma power was decreased and spontaneous gamma power was increased. Gamma coherence and phase locking value did not show significant changes. Cross-frequency coupling was shifted from gamma-delta to gamma-theta rhythm.

**Conclusion:** In summary, we found that acoustic stimulation at 40Hz can reduce Aβ in the brain and restore the gamma band oscillations and the frontoparietal connectivity. Our data suggest that acoustic stimulation might alter the natural deterioration processes of AD and have a therapeutic potential in AD.

## Introduction

Alzheimer’s disease (AD) is the most common cause of dementia ^1-3^ and understood as a complex disease that affects behavior and cognition through various pathophysiological mechanisms ^4^. The deposition of amyloid-beta (Aβ) peptide is one of the major pathological findings associated with AD ^5^. In AD, the balance between formation and clearance of Aβ is impaired ^6^. Under normal physiological conditions, glia cells, especially microglia and astrocytes, play crucial roles in maintaining the balance. The aggregated Aβ is degraded by microglia and astrocyte ^7^ and the soluble Aβ is eliminated through the perivascular pathway ^8-11^. In spite of recent failures of drug development targeting Aβ and skepticism, the ‘amyloid hypothesis’ are still one of the leading theory of AD pathogenesis ^12^.

Neuronal activity with gamma band (30-70 Hz) oscillations (GBO) in the human electroencephalogram (EEG) is known to be associated with human sensation, perception, cognitive process, such as information storage, retrieval, and integration ^13-16^. Cortical GBO are produced by the interplay between inhibitory interneurons and excitatory pyramidal neurons cellular ^17^. It is noteworthy that the patients with AD have shown abnormal GBO. Several studies have reported decreased activity of the spontaneous GBO in AD ^18,19^. The synchronization in spontaneous GBO was also suppressed compared to healthy controls ^19,20^. In contrast, evoked GBO as a steady-state response (SSR) to the repeated visual, auditory or somatosensory stimuli at a certain frequency increased in patients with AD ^21-23^.

The deposition of Aβ impairs synaptic plasticity at glutamatergic synapses, forms a synaptic depression with the morphological change of dendritic spine ^24,25^, and the GBO ^26^. Therefore, immunotherapy has been conducted aiming at the reduction of accumulated Aβ. Intracranial or systemic administration of immunotherapeutic agents reduced Aβ deposition by mechanisms both associated with and independent of microglial activation ^27,28^. Following studies have reported decreased Aβ plaques, increased soluble Aβ and altered shape of dystrophic neurite ^29-33^. However, immunotherapy did not improve the overall survival in phase 1 clinical trials with side effects ^34^.

The functional abnormalities of PV neuron and fast-spiking basket cells are origins of abnormal function of GBO ^35^. In mice with the mutated APP, damaged PV cells produce network dysfunction. Restoration of the interneuron-specific and PV cell predominant voltage-gated sodium channel (VGSC) subunit improved recovery of gamma oscillation and memory impairment and also resulted in improved survival outcome ^36^. However, this method is difficult to apply to clinical practice.

Recently, entrainment of the brain rhythm with GBO were used as a therapeutic method for AD ^26^. Optogenetic stimulation of PV neuron in the hippocampus and visual stimulation at 40 Hz using noninvasive flickering light also decreased Aβ peptides and increased activated microglia in the brain. However, their therapeutic effects were limited to hippocampus and visual cortex, respectively. To achieve a functional improvement, a broader area of the brain needs to be affected by the therapeutic effects. Given the necessity of other stimulation strategy, it is notable that rhythmic auditory stimulations can drive and entrain brain oscillations and this phenomenon is known as ‘auditory steady state response (ASSR)’ ^37^. Depending on the specific frequency stimulus, the auditory beat can induce or inhibit particular frequencies of the brain rhythms ^38^. In a clinical pilot study showed that rhythmic sensory stimulation at 40 Hz with sound stimuli improved cognition of AD patients ^39^. Therefore, we hypothesized that evoked GBO using 40 Hz acoustic stimulation (AS) can decrease Aβ and increase microglia and the neuropathological improvements may cause neurophysiological benefits, such as normalization of GBO and brain connectivity.

## Results

### Evoked gamma oscillation facilitates degradation of Aβ plaque with microglia activation

Aβ plaques were diminished in Tg+/Stim+, but not in Tg+/Stim-(Figure 1a). The number of Aβ plaque decreased in the pre-and infralimbic(PIL) and hippocampus(HC) regions (46.8% and 60.0% respectively), and it was significantly different between Tg-/Stim-and Tg+/Stim-, Tg-/Stim-and Tg+/Stim+ (Figure 1b, *p*<0.01, *p*<0.05).

**Fig. 1.**
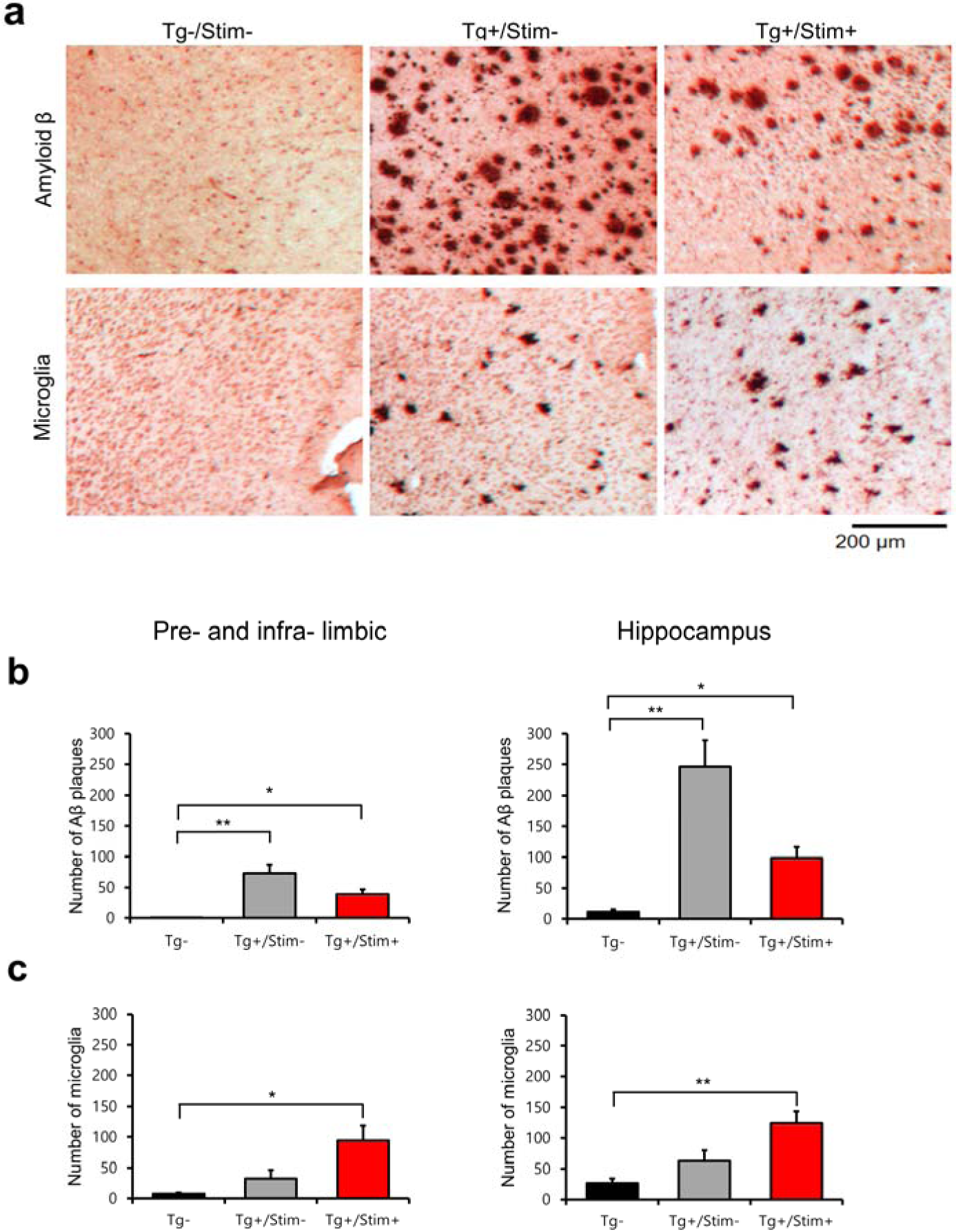
Reduced number of amyloid beta (Aβ) plaques in 5XFAD mice after acoustic stimulation at 40 Hz. a. Representative histologic findings with the immunohistochemistry for Aβ (*upper row*) and microglia (*bottom row*) in the pre-and infra-limbic cortex. **b, c** Quantification of the number of Aβ plaques and microglia in the pre-and infra-limbic cortex and hippocampus, respectively. Black, red, and gray bars indicate Tg-, Tg+/Stim+, and Tg+/Stim-, respectively. ^*^p < 0.05, ^**^p < 0.01 Mann-Whitney test following Kruskal-Wallis test or Dunnett T3, Bonferroni(equal variance assumed) post-hoc test following ANOVA test.

Microglia was aggregated in Tg+/Stim+ and Tg+/Stim-, but not in Tg-/Stim-(Figure 1a). There was a significant difference between microglia number per ROI in PIL and HC of Tg+/Stim+, Tg+/Stim-and Tg-/Stim-(Figure 1c, *p*<0.01). Microglia number in PIL and HC increased in Tg+/Stim+ than Tg-/Stim-showed statistical difference (Figure 1c, *p*<0.05, *p*<0.01). And, microglia number increased in PIL and HC (188.3% and 98.9%, respectively), showed no significant difference between Tg+/Stim-and Tg+/Stim-(Figure 1c, *p=*0.34, *p=*0.47). Evoked gamma oscillation facilitates microglia aggregation in Tg+/Stim+ than Tg-/Stim-.

### Evoked gamma oscillation decreased Aβ-40, 42 (soluble and insoluble) in PIL

There was a significant difference Aβ level of PIL between Tg+/Stim+, Tg+/Stim-and Tg-/Stim-(Figure 2, *p*<0.01). Tg+/Stim+, Tg+/Stim-have significantly increased Aβ-40, 42 (soluble and insoluble) levels than Tg-/Stim-. ELISA results show significant reduction of soluble Aβ-40 and -42 peptides (45.5% and 67.2%, respectively) in PIL in Tg+/Stim+ compared to Tg+/Stim-(Figure 2, *p*<0.05). On the other hand, there was no statistical difference in other levels of Aβ between Tg+/Stim+ and Tg+/Stim-.

**Fig. 2.**
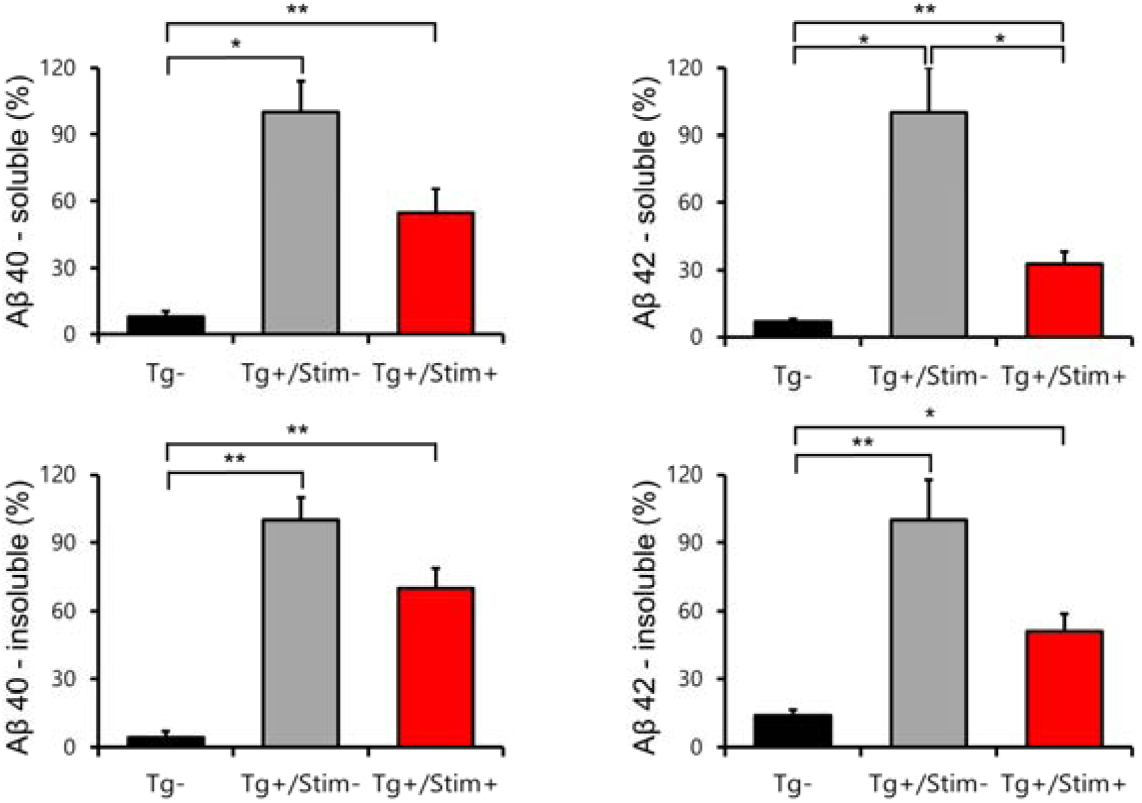
Reduced soluble Aβ40 and Aβ42 levels in pre-and infra-limbic after acoustic stimulation at 40 Hz. Black, red, and gray bars indicate Tg-, Tg+/Stim+, and Tg+/Stim-, respectively. ^*^p < 0.05, ^**^p < 0.01 Mann-Whitney test following Kruskal-Wallis test or *post-hoc* Dunnett T3 test following ANOVA test with welch correction.

### Gamma power in electroencephalography changed after 2 weeks of acoustic stimulation at 40 Hz

We measured the effect of 2 weeks of acoustic stimulation on local field potentials (LFP) to click-trains at gamma band frequency. Wavelet spectral analysis was performed on the averaged waveforms of evoked LFPs within the temporal window of stimulus. Figure 3 show the wavelet power spectral density and scale is corrected for inter-group comparison. Figure 3a show more increased evoked gamma band power on Tg+/Stim+ in day 1 than Tg-/Stim-. Figure 3a also show the reduction of evoked gamma power in Tg+/Stim+ in day 14 than Tg+/Stim-. The data were normalized by its pre-stimulus gamma band power and fold changes were analyzed. Exaggerated evoked gamma power was observed in Tg+/Stim+ in day 1 or Tg+/Stim-(Figure 2b, c). We performed repeated measure analysis of variance (RM ANOVA) for serial follow up with evoked or resting gamma power of Tg+/Stim+ at different time point. Tg+/Stim+ show statistically no significant reduction in evoked gamma power among day 1, 7 and 14 (*p* = 0.57). Tg+/Stim+ significantly increased resting-spontaneous gamma power (RS-gamma power) in day 7 than day 1, day 14 than day 1 (*p* < 0.05, *p* < 0.05).

**Fig. 3.**
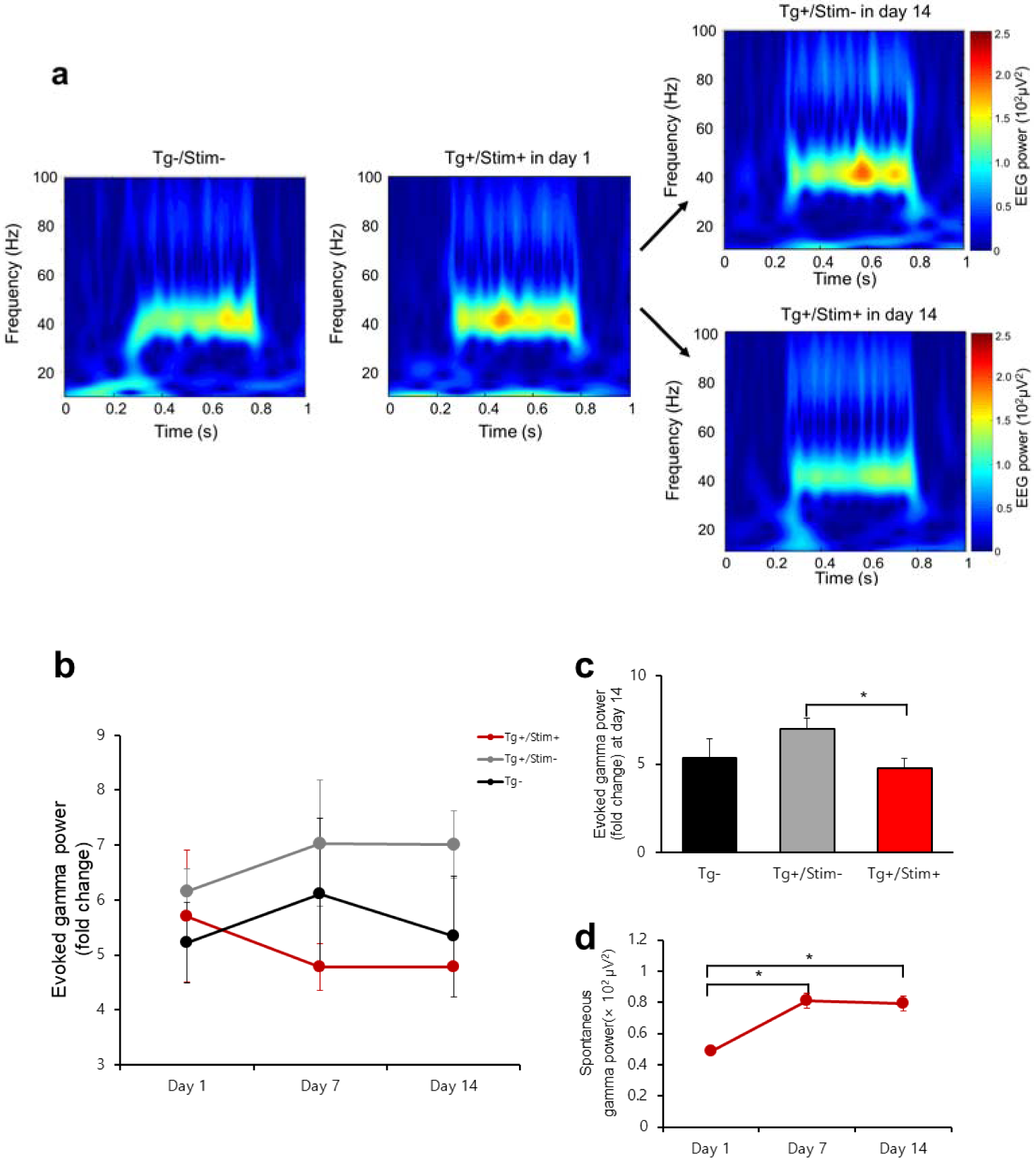
Evoked and spontaneous gamma power in electroencephalography changed after 2 weeks of acoustic stimulation at 40 Hz. a. Time-frequency analysis of the grand averaged EEG data from the auditory steady-state response (ASSR) at 40Hz using the wavelet transformation. **b** Time course of the evoked gamma power in Tg-, Tg+/Stim-, and Tg+/Stim+ groups at the stimulation day 1, 7, and 14. Tg+/Stim+ was not statistically significant by repeated measure ANOVA. **c** The evoked gamma power at day 14 in Tg-, Tg+/Stim-, and Tg+/Stim+ groups, depicted as black, gray and red bars, respectively. There is statistical difference between Tg+/Stim-and Tg+/Stim+. ^*^P<0.05 by Mann-Whitney test. **d** Time course of resting gamma power in Tg+/Stim+ group.^*^ P<0.05 by Bonferroni correction following repeated measure ANOVA.

### Connectivity between frontal and parietal electroencephalography changed after 2 weeks of acoustic stimulation at 40 Hz

We performed a two-way RM ANOVA for serial follow up with FP connectivity at different groups and time points. FP connectivity was calculated as coherence and phase locking value. Coherence is calculated at evoked gamma duration, and phase locking value(PLV) is calculated at RS gamma period. Figure 4a showed no statistically significant difference in coherence during gamma test of day 1, 7, and 14 (*p =* 0.108). Tg+/Stim+ showed slight decrement from day 1 to 7, and increment in day 14 (0.82±0.02, 0.73±0.03, 0.76±0.03); on the other hand, Tg+/Stim-showed stable value on day 1, 7, and 14 (0.79±0.03, 0.79±0.04, 0.78±0.03). Tg-/Stim-showed slight reduction in day 1, 7, or 14 (0.62±0.10, 0.56±0.06, 0.57±0.09). Figure 4b showed no statistically significant difference of PLV. Tg+/Stim+ showed increment on day 1, 7, and 14 (0.5±0.1, 0.52±0.11, 0.59±0.07); on the other hand, Tg+/Stim-showed a serial reduction of resting spontaneous gamma phase locking value on day 1, 7, and 14 (0.63±0.03, 0.63±0.06, 0.43±0.15). Tg-/Stim-showed a slight change in phase locking value in day 1, 7, or 14 (0.65±0.05, 0.65±0.1, 0.68±0.08).

**Fig. 4.**
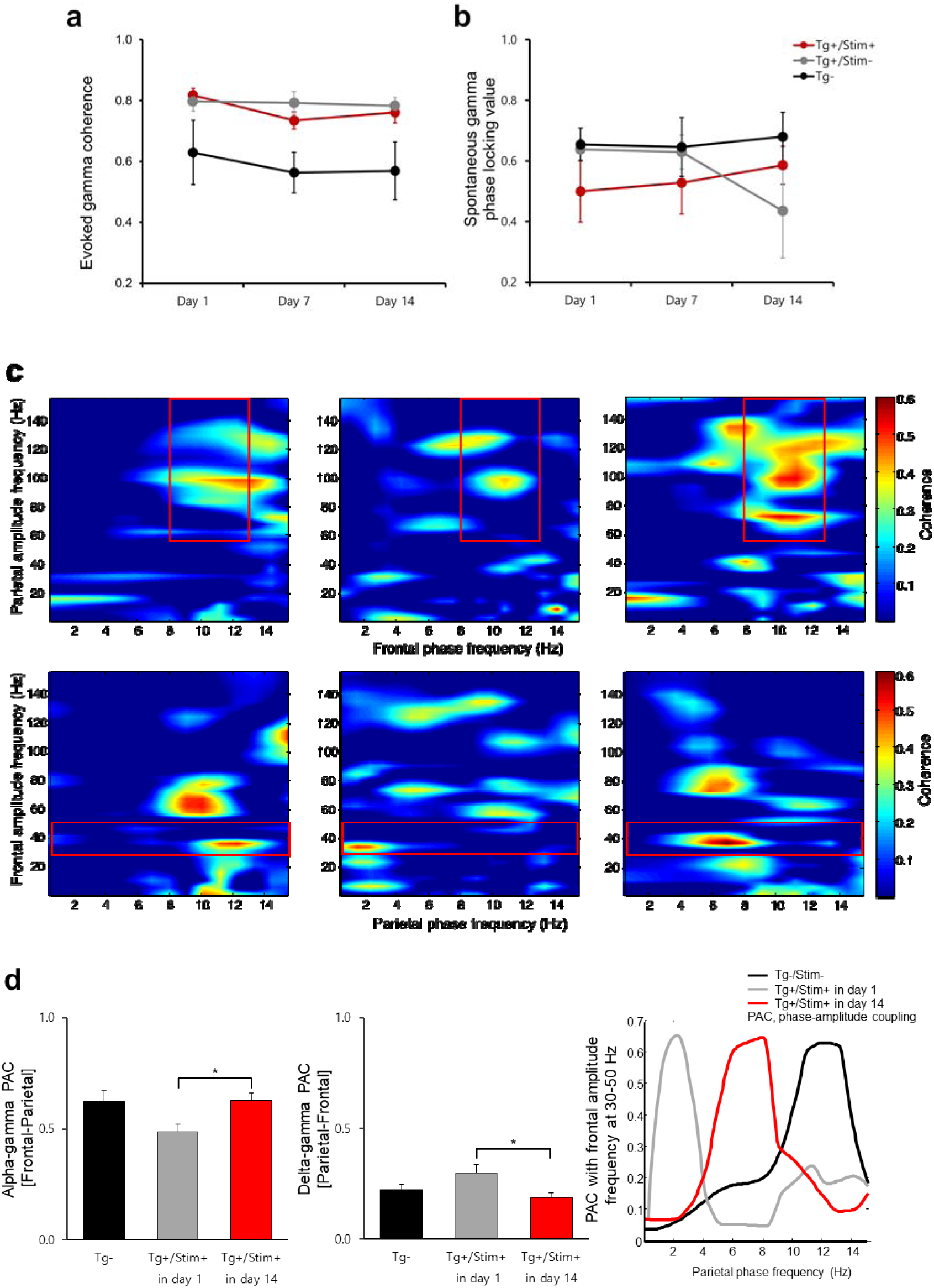
Connectivity between frontal and parietal electroencephalography changed after 2 weeks of acoustic stimulation at 40 Hz. **a, b** Time course of the evoked gamma coherence and the resting gamma phase locking value. Tg-, Tg+/Stim-, and Tg+/Stim+ groups were depicted as black, gray and red circles and lines, respectively. Tg+/Stim+ was not statistically significant by repeated measure ANOVA. **c** Cross-frequency phase-amplitude coupling (PAC) of the frontal and parietal grand-averaged EEG during the auditory steady-state response tests. Parietal amplitude frequency and frontal phase frequency coupling (*upper row*) and frontal amplitude frequency and parietal phase frequency coupling (*bottom row*) were plotted. The color bar indicates coherence (ranging from 0 to 1), with no coherence (0) to perfect coherence (1). Red rectangles represent the area of interest (ROI). **d** Quantification of the maximal PAC values in the ROIs. Black, gray and red bars represent Tg-, Tg+/Stim+ at baseline, and Tg+/Stim+ at day 14, respectively.^*^ P<0.05 by *post-hoc* Bonferroni test following ANOVA.

Phase-amplitude coupling (PAC) changes of the frontal and parietal region in Tg-/Stim-and Tg+/Stim+ in day 1, 14 are observed in Figure 4c, d. PAC patterns including high-gamma activity are analyzed. Top and bottom layer showed that grand-averaged cross-frequency PAC in Tg-/Stim-and Tg+/Stim+ in day 1, 14. PAC is performed as the analysis amplitude (0-160 Hz) and analysis phase (1-16 Hz) combination for F-phase / P-amplitude (top row) and P-phase / F-amplitude (bottom row). The emphasis is that there is low coupling between alpha-phase(F) and high gamma amplitude(P) in Tg+/Stim+ in day 1 (middle column) compared to the Tg-/Stim-(left column) and Tg+/Stim+ in day 14 (right column) (areas highlighted in red square). Delta-phase(P) and evoked gamma-amplitude(F) coupling is exaggerated (areas highlighted in red square) in Tg+/Stim+ in day 1 compared to the Tg-/Stim-and Tg+/Stim+ in day 14. Comparison of maximum PAC values which contain analysis amplitude (30-160 Hz) and alpha phase (8-13 Hz) combination for F-phase / P-amplitude is statistically significant, and comparison of maximum PAC values which contain analysis amplitude (30-160 Hz) and delta phase (1-4 Hz) combination for P-phase / F-amplitude is statistically significant between Tg+/Stim+ in day 1 and day 14 (Figure 4d, *p* < 0.05, *p* < 0.05). Mainly coupled frequency of parietal phase with 40Hz frequency of frontal amplitude is high alpha(11-13 Hz) in Tg-/Stim-. That is delta (0-4 Hz) in Tg+/Stim+ in day 1, and is shifted to theta (4-8 Hz).

## Discussion

The AD is characterized by progressive deposition of Aβ and tau protein, resulting in abnormalities of GBO and altered synchronization problems ^2,18,40^. Our results showed that 2 weeks of GBO entrainment by acoustic stimulation (AS) at 40Hz facilitated degradation of Aβ plaque with microglia activation. The effects were shown in both PIL cortex and HC which are important areas of the brain for the cognitive function such as executive function and memory. We also found that the evoked GBO were decreased, whereas spontaneous GBO were increased through the stimulation period. Frontal gamma rhythms are coupled with parietal delta at baseline, but the coupling of frontal gamma shifted to parietal theta rhythms after acoustic stimulation.

The number of Aβ plaques in PIL and HC reduced in Tg+/Stim+ than Tg+/Stim-with increased number of microglia. *Post hoc* pairwise comparisons of the number of microglia showed statistical significance only between acoustic stimulation group and wild type, but not between with and without acoustic stimulation. Tg+ mice without acoustic stimulation also showed slight increases of the microglia as a physiological response to increased Aβ, which might reduce the statistical significance. We also performed 2-minute ASSR tests at 40 Hz in day 1, 7 and 14 to measure the evoked GBO responses, which may have caused partial acoustic stimulation effects.

AD has a strong genetic characteristics, and recent observation in genome technology report missense variants in triggering receptor expressed on myeloid cells-2 (TREM2) are correlated with a 2-4 fold increased risk of developing AD ^41,42^. TREM2 is essential for Aβ-related microglial responses in mouse models and human. Though the mechanistic basis for reduced microglia is unclear, TREM2 deficiency may cause reduction of microglial response with increased apoptosis, reduced proliferation, or infiltration of peripheral macrophages ^43^. TREM2 mediates early microglial response, and limits diffusion and toxicity of amyloid plaques ^44^. Our results showed that acoustic stimulation at 40 Hz enhanced the microglial response and reduced Aβ deposition. Further researches are warranted to determine whether the effects of acoustic stimulation is mediated by TREM2.

We compared our results with previous studies about the recovery of GBO ^26,39^. In the study by Iaccarino *et al.* soluble and insoluble Aβ decreased in the target region. Although the optogenetic stimulation and visual flickering methods differ from acoustic stimulation, the common denominator between the two stimuli are the evocation of GBO. In Tg+/Stim+ and Tg+/Stim-, the level of soluble and insoluble Aβ was higher than Tg-/Stim-. The level of soluble Aβ-42 was significantly lower in Tg+/Stim+ than Tg+/Stim-, while the level of insoluble Aβ was not statistically significant. The reason why the soluble Aβ decrease significantly, and insoluble Aβ decrease did not show the statistical difference in our results might be the cerebral amyloid angiopathy (CAA) involving capillaries in the progression of the AD ^8^. Similarly, when insoluble Aβ was degraded by immunotherapy via activated microglia, CAA might be aggravated. Consequently, inadequate drainage of soluble/insoluble Aβ led to increased micro-hemorrhage and decreased survival outcome ^8,29-31^.

Considering the characteristics of AD mice, in which Aβ deposition is observed from the age of 2 months ^45^, it is possible that drainage of insoluble Aβ was not performed correctly with progressed CAA due to severe symptoms in 6-month-old mice ^46-49^ used in this experiment. Although insoluble Aβ might have been degraded to soluble Aβ by microglial activation, insoluble Aβ may remain unaltered in case of progressed CAA ^8,29-31^. On the other hand, because 3-month-old mice may not have progressed CAA, the drainage of Aβ would have been intact and thus both soluble and insoluble Aβ decreased in the study of Iaccarino *et al*.

These differences suggest that therapeutic methods through degradation of Aβ plaques in patients with progressed dementia should consider the timing of therapeutic point, and the drain pathway of soluble Aβ. Aging is characterized by CSF compartment in brain parenchyma due to cerebral atrophy and decreased cerebrospinal fluid turnover ^50^. Choroidal epithelium and ependyma degenerate in aging and also AD. 40% of patients with adult chronic hydrocephalus contain deposition of Aβ ^51^. With evidence of decreased CSF Aβ levels have been reported after an internal CSF shunt ^52^, we suggest a novel combination method of creating CSF shunt for drainage pathway during 40 Hz AS to treat progressed AD patients.

Verrett *et al*. reported the restoration of the GBO after correcting the abnormalities of the VGSC of PV cell followed by the recovery of cognitive function and the reduction of mortality in the AD animal model ^36^. In our study, we found that evoked GBO was reduced and the spontaneous GBO was increased, which might be neurophysiological evidence for the functional improvement after acoustic stimulation.

Altered N-methyl-D-aspartate receptors (NMDARs) leads to abnormal function of inhibitory GABAergic interneurons, especially those containing PV interneurons, and the ultimate result is disrupted inhibition of excitatory pyramidal neurons ^53,54^. Considering excessive levels of glutamate result in neurotoxicity, NMDAR has been suggested to be a therapeutic target in neurodegenerative disease ^54^. Because over-activation of NMDARs cause neurotoxicity, partial NMDAR antagonist (called memantine) blocks the NMDA glutamate receptors to normalize the glutamatergic system and improve cognitive and memory deficits ^55^. NMDAR antagonist enhanced resting spontaneous broadband gamma activity ^56,57^. We found that spontaneous GBO in Tg+/Stim+ mice which received daily acoustic stimulation for 2 weeks was increased at day 7 and 14. The increase in resting spontaneous GBO is also reported in previous studies to show the therapeutic effectiveness. ^18,19,36^.

We also found the overactivity of GBO evoked by acoustic stimulation in 5XFAD mice. These results are consistent with the previous studies. Measured activities of evoked gamma band power in AD patients was increased compared to the normal group ^23,40^. These results are associated with studies that analyzed P50 using a double-click paradigm or auditory steady-state paradigm known as a measure of inhibition of the cortex ^21,22^. Several studies using this paradigm in AD patients showed an increase in the amplitude of the second click sound ^21^. Similarly, normal controls with a family history of the AD have similar results, suggesting that impaired cortical inhibition may occur very early in the course of AD disease ^58^. It is notable that auditory stimulation and response play an essential role in the early diagnosis. A comparative study with patients with mild cognitive impairment (MCI), which is likely to progress AD, as well as patients with the AD, suggests that disorders with abnormal which GBO lead to cognitive impairment were reported ^40^. Interestingly, after 14 days of acoustic stimulation, we found a reduction of evoked GBO in Tg+/Stim+ mice than the age-matched Tg+/Stim-mice.

Neural synchronization between the frontal and parietal electrodes in 5XFAD mice was analyzed. We only observed trends of increased coherence and decreased phase locking value in 5XFAD, but there was no statistical significance. Theoretically, there is a possibility that the coherence at the gamma frequency increase and the phase locking value decrease in the Tg+/Stim-. For the computation of coherence, both amplitude and frequency of the EEG are utilized. Thus, when the amplitude is large at a certain frequency, the weight becomes larger in the calculation process than other frequencies. Therefore, coherence increases when the acoustic stimulation-evoked gamma band power is increased. On the other hand, in the case of the instantaneous PLV, it may be effective to observe the functional decrease of the connectivity because only the phase excluding amplitude and frequency is considered. Previous studies have been reported increased coherence in treated AD patients ^59,60^, and to decrease PLV and global field synchronization in the progression of AD ^18-20^. We showed time courses of the decreasing pattern of PLV in Tg+/Stim-mice in comparison with Tg+/Stim+ mice. Two weeks of acoustic stimulation might have contributed to a sustained pattern of PLV, whereas a progressive reduction of PLV was shown for the Tg+/Stim-mice.

Cross frequency phase amplitude coupling (CF-PAC) emphasized that there is lower coupling between alpha-phase(F, frontal) and high gamma amplitude(P, parietal) in Tg+/Stim+ in day 1 compared to wild type mice or Tg+/Stim+ in day 14. During cognitive processing, high gamma CF-PAC is important for effective interregional and local communication ^9,26^. Alpha-phase high-gamma coupling in wild type mice was suggestive about effective bidirectional modulation from frontal to auditory areas. The reduction of PAC of Tg+ mice in day 1 implies that frontal top-down and parietal bottom-up control may be impaired. This result showed that attention resources could not be appropriately processed. The prefrontal associative cortex is responsible for what is called "top-down" attention, based on the relevance of current tasks to internal goals ^40,53,54^. After 2 weeks of acoustic stimulation, PAC between alpha-gamma coupling is increased significantly to the level of wild type mice. Gamma amplitude (F) is coupled with alpha frequency (P) in wild type mice but Tg+ mice showed the strongest coupling between gamma (F) and delta (P). After 2 weeks of acoustic stimulation, the PAC is shifted from gamma (F) – delta (P) to gamma (F) – theta (P). Therefore, acoustic stimulation seemed to cause changes in cross-frequency coupling, which might reflect the improved information processing and cognitive function.

To our best knowledge, this is the first study to confirm the therapeutic effects of acoustic stimulation for GBO entrainment in a mouse model of AD with pathological validation and serial changes of EEG characteristics including GBO, coherence, PLV and CF-PAC. We discovered that 2 weeks of acoustic stimulation at 40 Hz can modify the pathophysiology of Aβ deposition in the cortex of 5XFAD mice, and restoration of GBO and cross-frequency coupling. Further research with acoustic intervention for AD is warranted.

## Materials and methods

### 2.1 Animals

Adult (5-month-old) male 5XFAD mice were prepared. The 5XFAD mouse model (Tg6799, Stock 6554) was purchased from Jackson Laboratory. This model is a double transgenic that overexpresses not only PS1 M146L and L286V but also APP Swedish, Florida and London mutations, which are under the control of the murine Thy-1-promoter. Swedish mutation produces higher levels of total Aβ; on the other hand, Florida, London, M146L, and L286V mutations characteristically increase the production of Aβ_x-42_. Mice of the B6SJLF1 / J background strain have high APP expression correlated with accelerated accumulation of Aβ_x-42_ in comparison to Aβ_x-40_. 21 males were used in this experiment, and all mice were handled with Gwangju Institute Science and Technology (GIST) Laboratory Animal Resource Center (LARC) guidelines for animal care. Mice were bred and kept in a 12/12 hour light-dark cycle (light on at 9 a.m.) a temperature and humidity controlled room at 20°C ± 2°C, 55%±5%. Water and food were freely available *ad libitum*. All animal procedures have been approved by the ethics committee of GIST LARC which fulfilled with Association for Assessment and Accreditation of Laboratory Animal Care International guidelines. 15 males were 5XFAD mice, and 6 males were their wild-type littermates. 3 of 15 5XFAD males underwent behavioral tests before sound stimulation and after 2 weeks of sound stimulation. 12 of 15 5XFAD males were randomly separated into 2 groups. Group 1 [Tg(+)(ASSR(+)) consisted of 7 mice stimulated for 2 hours per day for 2 weeks. Group 2 [Tg(+)ASSR(-)] consisted of 5 mice and group 3 [wild-type littermates] consisted of 6 mice that simply received gamma test at day 0, 7, and 14.

### 2.2 Surgical preparation, electrode implantation

The animal was anesthetized with 4% isoflurane and maintained with 0.5 to 1.5% isoflurane in a stereotaxic frame. 0.1mg/kg of ketofen was injected subcutaneously before surgery to control pain. The blanket was used to maintain body temperature. Mouse eye was covered with protecting ointment. After shaving incision line, incision of the midline using scalpel and the cranium was exposed, five holes of 0.8mm were drilled over the frontal (AP: 1.0 mm, ML: 1.0 mm), parietal (AP: -3.5 mm, ML: 1.0 mm), ground (AP: -2.0 mm, ML: 1.5 mm), anchor (AP: -2.0 mm, ML: -1.5 mm), and reference (AP: -5.3 mm, ML: 0.0 mm). EEG electrodes of screw type were inserted at a depth that would hardly touch or touch the brain in drilled holes, and EMG electrodes were inserted in neck muscles. 1×2mm screws were inserted as an anchor for a metal holder that was cemented to the skull with dental cement. The screws with wire were fixed with dental cement, and after the cement had completely hardened, the remnant parts of wires and the connector (PINNACLE Technology Inc, Oregon, US) were fixed on the vertex of the skull by soldering and dental cement. In the end, the simple suture was performed after wound irrigation. Animals were then housed in a cage for more than 1 week for postoperative recovery.

### 2.3 In Vivo EEG/EMG Recordings

Continuous EEG/EMG (8200-K1-SL amplifier; Pinnacle Technology) were recorded before, during, and after 40 Hz auditory stimulation. We were careful about circadian timing and performed 2-hour auditory stimulation and recorded EEG from 22:00 (ZT 13) to 00:00 (ZT 15). This ensured that the animals were in their physiologic activity, and reduced the period of falling asleep during a stimulation session. Sleep was monitored with EEG/EMG signals. The EEG/EMG signal (2 KHz sampling, 100 Hz lowpass filtered; Pinnacle PAL8200/Sirenia software) response to auditory stimulation was recorded using WinWCP software (Strathclyde Institute of Pharmacy and Biomedical Sciences).

### 2.4 Auditory stimulus

Auditory clicks (20 rectangular pulses of 10ms duration per 500ms, 40Hz) per sec were generated by custom-built programs under WinWCP (Strathclyde Institute of Pharmacy and Biomedical Sciences) condition and delivered via a speaker, which created the sound pressure of 90dB in the soundproof room. There was a stimulation time of 500ms and a resting time of 700ms with a cycle of 1.2 seconds. The resting time consisted of the system time for the next recording of 200ms and the recording time, which is consisted of 250ms before stimulation and 250ms after stimulation. Recording and click trains were 1 cycle/sec, and each train had 200ms interval. In the end, the mice received 6000 cycles of the train for 2 hours every day. We performed gamma tests at 0, 7, 14 days of an experiment to check the gamma response between the groups. The gamma test is repeated 200 cycles of the above 1.2 seconds cycle, which takes a total of 4 minutes.

### 2.5 EEG Data Analysis

Offline processing was performed in the customized code for Matlab. Wavelet transformation (WT; continuous wavelet from Matlab Wavelet Toolbox; frequencies represented from 1 to 100 Hz) was performed. WT power was measured from 38 to 42 Hz gamma band.

We calculated magnitude-squared coherence between frontal and parietal epidural electrode. Using mscohere function in MatLab, we obtained sequences frontal and parietal EEG before and after stimulation for inspecting connectivity changes between frontal and parietal region. We also calculated phase locking value (PLV). For calculation of PLVs, the length of the time window that is required to obtain instantaneous phase information should be determined. We defined this window at resting spontaneous (pre-stimulation) period into 230ms lengths and stimulation period into 460ms lengths. Before the Hilbert transform, the frequency of interest should be determined. Local field potentials were filtered using a second order bandpass digital Butterworth filter with a higher and lower cut off adjusted to 2 Hz above and below the stimulus frequency. For example, lower and upper corner frequencies of 38 and 42 Hz respectively were used for the 40-Hz stimulus. PLV is defined as an average of the differences of instantaneous phase between the two signals. The PLVs have a range of values from 0 to 1, that indicates no phase locking to complete phase locking. Averaged evoked power, frontal-parietal coherence, phase-locking data were quantified. Gamma band wavelet power expressed evoked data relative to the resting spontaneous(pre-stimulus) data or resting spontaneous data itself.

Cross-frequency coupling (CFC) between frontal and parietal electrode was evaluated using the PAC values. We calculated the coherence between a low-frequency signal and the time-course of the power at the higher frequency. Signals containing phase was at 1 to 16 Hz, and signals containing amplitude was at 30 to 160 Hz. PAC values were calculated for each frequency pair for LFP recorded during auditory stimulation. The PAC analysis was conducted on the 60 Hz notch-filtered EEG data. To generate color-maps shown in the figure, we use grand averaged EEG in a particular group (Tg-/Stim-, Tg+/Stim-and Tg+/Stim+). To calculate spectrograms of EEG activity averaged at low-frequency phase troughs with the highest PAC, signals were first filtered in the delta band using a narrow band-pass filter (a symmetrical finite impulse response filter with a specific nesting frequency within the delta frequency range [1-4 Hz]), and then amplitude troughs were identified from the filtered signals. Using Brainstorm software, a time-frequency decomposition was performed to obtain the power of all averaged time-frequency plots. Alpha (8-13 Hz) phase and gamma (40-160 Hz) amplitude cross-frequency coupling were investigated based on papers that the phase of lower frequency oscillations modulate the amplitude of gamma oscillations ^61,62^. The maximum PAC values of these frequency ranges were analyzed among Tg-/Stim-, Tg+/Stim+ in day 1 and 14.

### 2.6 Immunoassay and immunohistochemistry

## Mouse brain homogenization

Mice were deeply anesthetized by intraperitoneal injection of Zoletil (Virbac, SA, Carros, France), and then transcardiacally perfused with filtered PBS. Brains were removed, dissected, frozen in dry-ice cold isopentane (Sigma-Aldrich, St. Louis, MO, USA), and stored at -80 °C until use. Medial prefrontal cortex (mPFC) was extracted as 1:4 (4 µl/mg of brain) of ice-cold TBS extraction buffer (50 mM Tris-buffered saline, pH 7.6) containing complete protease inhibitor cocktail (Roche Diagnostics) using a tissue grinder with Teflon pestle on ice. The homogenates were centrifuged at a speed of 200,000 g for 20 min at 4°C (Beckman Optima TL Ultracentrifuge, TLA100.4 rotor). The supernatants (TBS-soluble fraction) were kept at -80 °C before the assay. After adding 4 volumes (based on initial hemisphere weight) of 6M GuHCl solution (6M GuHCl, 50 mM Tris-HCl, protease inhibitor cocktail, pH 7.6), samples were incubated in 6M GuHCl solution for 1 hr at 25°C and finally centrifuged at a speed 200,000 g for 20 min at 4°C to extract GuHCl-soluble fraction.

## Sandwich Enzyme linked immunesorbent assay (ELISA)

Levels of amyloid beta 1-40 and 1-42 in the brain extracts were assayed with sandwich ELISA kit (#KHB3481 for amyloid beta 1-40, #KHB3441 for amyloid beta 1-42, Thermo Scientific) according to the manufacturer’s guidance. Optical densities (at 450 nm) were measured by Tecan Infinite F50 microplate reader (TECAN, Austria). Assay sensitivity was <6 pg/ml for amyloid beta 1-40 and <10 pg/ml for amyloid beta 1-42.

## Immunohistochemistry

Brains were fixed in 4% paraformaldehyde solution for overnight, and then placed in 30% sucrose solution at 4 °C until they sink. Serial coronal sections (30-µm thickness) of the whole brain were obtained and were stored in cryoprotectant solution (30% RNase free sucrose, 30% ethylene glycol, and 1% polyvinyl pyrrolidine in 100 mM, pH 7.4) at -20 °C. All immunohistochemical stainings were executed using the free-floating method as described below. Samples were incubated in 70% formic acid solution for 10 min, washed, and then incubated in 1% H 2 O 2 in 0.1 M PBS (pH 7.4) for 10 min to block endogenous peroxidase activity. Tissues were reacted in a buffer containing 0.2% Triton X-100 (PBS-T), 10% normal goat serum (NGS) for 1 h. Sections were then incubated for 3 days at 4°C with a mouse anti-amyloid beta 1-42 (1:1,000; Biolegend) or rabbit anti-iba-1 antibody (1:1000; Wako Pure Chemical Industries, Osaka, Japan, #019-19741). After washing, they were incubated with biotinylated goat anti-mouse IgG (1:1,000; Vector Laboratories, Burlingame, CA, USA) or biotinylated goat anti-rabbit IgG (1:1,000; Vector Laboratories) for 12–24 h at 4 °C. After several rinses with PBS-T, the sections were incubated in an avidin–biotin– peroxidase complex (1:250; Vector Laboratories) for 1 h. After rinsing, sections were incubated for 10 min in 0.05% (w/v) diaminobenzidine solution containing 0.3% H 2 O 2.

The sections were then washed with PBS, mounted on glass slides, dried overnight, dehydrated, and cover-slipped under Permount (Fisher Scientific, Fair Lawn, NJ, USA). Images showing immunoreactivites were captured using a ProGress C14 camera (Jenoptik,Jena, Germany) mounted on a Zeiss axioscope microscope (Zeiss, Germany).

### 2.7 Statistics

Normal distribution of data was validated with Kolmogorov-Smirnov test and Shapiro-Wilk test. For group comparison at a separate time point, we used the one-way ANOVA followed by the Bonferroni multiple comparisons for normally distributed data and Kruskal-Walis and Mann-Whitney Rank Sum Test for non-normally distributed data. Serial data was analyzed with repeated measure analysis of variance (RM ANOVA). The interaction between variables was also calculated. All data are shown as an average ± standard error of the means (S.E.M). The statistical significance levels are indicated by one or two asterisks in figures (*p<0.05,**p<0.01).

